# Cold-induced hyperphagia requires AgRP neuron activation in mice

**DOI:** 10.1101/2020.05.21.107896

**Authors:** Jennifer D. Deem, Chelsea L. Faber, Christian Pedersen, Bao Anh Phan, Sarah A. Larsen, Kayoko Ogimoto, Jarrell T. Nelson, Vincent Damian, Megan A. Tran, Richard D. Palmiter, Karl J. Kaiyala, Jarrad M. Scarlett, Michael R. Bruchas, Michael W. Schwartz, Gregory J. Morton

## Abstract

To maintain energy homeostasis during cold exposure, the increased energy demands of thermogenesis must be counterbalanced by increased energy intake. To investigate the neurobiological mechanisms underlying this cold-induced hyperphagia, we asked whether agouti-related peptide (AgRP) neurons are activated when animals are placed in a cold environment and, if so, whether this response is required for the associated hyperphagia. We report that AgRP-neuron activation occurs rapidly upon acute cold exposure, as do increases of both energy expenditure and energy intake, suggesting the mere perception of cold is sufficient to engage each of these responses. We further report that silencing of AgRP neurons selectively blocks the effect of cold exposure to increase food intake. Together, these findings establish a physiologically important role for AgRP neurons in the hyperphagic response to cold exposure.

## INTRODUCTION

In homeothermic species, maintenance of euthermia in the face of a wide range of ambient temperatures is critical for survival. In small homeotherms, countering cold stress requires not only that adaptive adjustments of heat production occur rapidly and potently, but that they are achieved without depleting body fuel stored in the form of fat (Gordon, 1993). Thus, when animals are housed in a cool environment, energy expenditure increases rapidly and markedly to generate the heat needed to maintain core body temperature (Cannon & Nedergaard, 2010), and a compensatory hyperphagic response prevents changes to body fat mass (Kaiyala, et al., 2015; Ravussin, et al., 2014). While much is known regarding the neurocircuitry underlying cold-induced thermogenesis (Madden, et al., 2019; Nakamura, et al., 2011; Tan, et al., 2018), the origins of cold-induced hyperphagia remain poorly understood.

One potential explanation for cold-induced hyperphagia is that it is mounted as a secondary response to the negative energy-balance state that results from cold-induced thermogenesis, analogous to what occurs in other states of negative energy balance. During caloric restriction, for example, the imposed negative energy balance and the associated reduction of body fat stores drives activation of adaptive homeostatic responses aimed at returning body fat stores to pre-intervention values. Reduced energy expenditure and increased food-intake drive are major components of this adaptive response, and both are thought to be primarily driven by humoral signals indicative of a negative energy state (e.g., reductions of leptin and insulin) (Schwartz, et al., 2000). It has, therefore, been reasonably assumed that the hyperphagic response to cold requires the same negative energy-state signals, and this hypothesis is supported by studies that have been conducted over time periods sufficient to induce hormonal changes (Bing, et al., 1998; Hardie, et al., 1996; Puerta, et al., 2002).

Among central targets of these humoral feedback signals are neurons that express agouti-related peptide (AgRP) located in the hypothalamic arcuate nucleus (ARC) (Hahn, et al., 1998) which, when activated, potently stimulate feeding (Aponte, et al., 2010; Atasoy, et al., 2012; Krashes, et al., 2011). However, recent evidence suggests that AgRP neurons are not regulated solely by humoral signals (Hahn, et al., 1998; Schwartz, et al., 2000; Zimmer, et al., 2019), but also by feed-forward mechanisms (Chen, et al., 2016; Lowell, et al., 2019) involving neurocircuits that integrate sensory cues from the environment (Betley, et al., 2015; Chen, et al., 2015, Mandelblat-Cerf, et al., 2015) such that a negative energy state may be prevented. Based on these observations, we hypothesized a role for AgRP neuron activation in the adaptive increase of food intake induced by cold exposure.

## RESULTS

### Effect of chronic cold exposure on determinants of energy balance and hypothalamic neuropeptide expression

Our first goal was to confirm previous findings regarding the effect of chronic cold exposure on energy homeostasis in normal mice. Using a mild, chronic cold-exposure paradigm (14°C for 5 days) and comparing outcomes to mice housed at room temperature (22°C) (**Figure 1A**), we found that, as expected (Cannon & Nedergaard, 2004; Morrison, et al., 2014), core body temperature did not change significantly relative to controls housed at room temperature (**Figure 1B**), presumably owing to an associated increase of heat production (**Figure 1C**). This thermogenic response was accompanied by a proportionate increase of energy intake (Kaiyala, et al., 2015; Ravussin, et al., 2014; Vallerand, et al., 1986) (**Figure 1D**), such that neither body weight nor body fat mass changed significantly over the course of the study (**Figure 1E-G**). Moreover, neither respiratory quotient (RQ) nor ambulatory activity changed significantly (data not shown).

**Figure 1.**
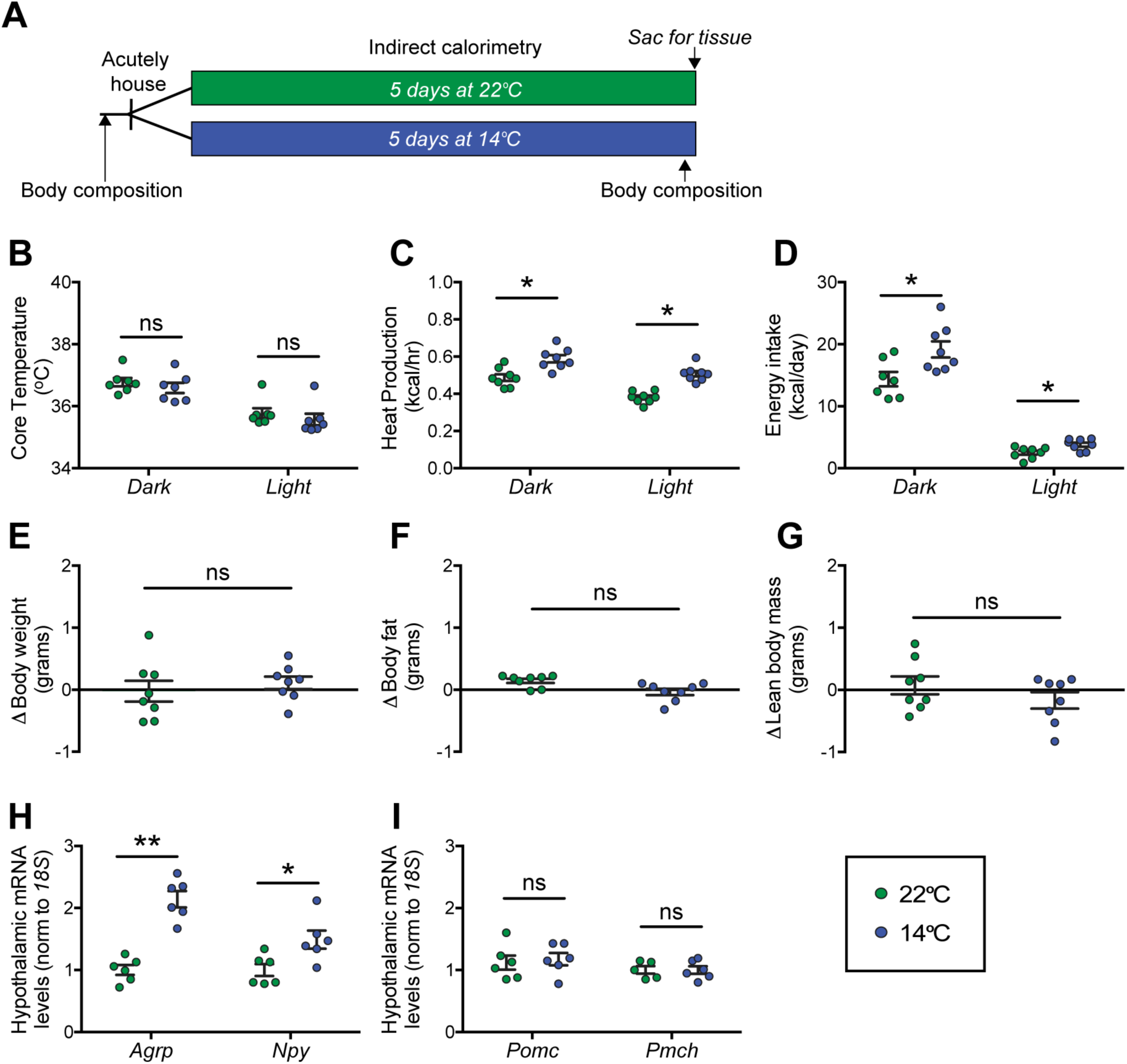
Effect of chronic cold exposure on determinants of energy balance and hypothalamic neuropeptide gene expression. (**A**) Adult male, wild-type mice were housed at the start of the dark cycle following body-composition measures and were maintained in temperature-controlled chambers set to either mild cold (14°C) or, as control, room temperature (22°C) for 5 days. Following a final body-composition analysis, animals were sacrificed and hypothalamic punches rapidly dissected. (**B**) Mean dark and light cycle core body temperature, (**C**) heat production, (**D**) and energy intake over 5 days while housed at either 22°C or 14°C. (**E**) Change in body weight, (**F**) body fat (**G**) and lean body mass at study end in the same mice, n= 7-8/group, mean ± SEM. Student’s t-test, *p<0.05 vs. 22°C. Hypothalamic mRNA levels of (**H**) agouti-related peptide (*Agrp*) and neuropeptide Y (*Npy*), (**I**) pro-opiomelanocortin (*Pomc*) and pro-melanin concentrating hormone (*Pmch*) were determined by qRT-PCR, n= 6/group, mean ± SEM. Student’s t-test, *p<0.05, **p<0.01 vs. 22°C.

At the end of the 5-day study, animals were sacrificed and hypothalami were rapidly dissected for RNA isolation. To identify candidate hypothalamic mediators of this cold-induced hyperphagic response, we used quantitative real-time PCR (qRT-PCR) to measure select neuropeptide transcripts in isolated hypothalamic RNA. Despite our use of a mild cold-exposure paradigm, consistent with previous work using stronger cold-exposure paradigms (∼4-5°C) (McCarthy, et al., 1993; Tang, et al., 2009), we found increased expression of orexigenic *Agrp* and *Npy* mRNA in animals housed at 14°C when compared to controls housed at room temperature (**Figure 1H**), while expression of *Pomc* and *Pmch* remained unchanged (**Figure 1I**). This evidence of enhanced *Agrp* mRNA expression cannot be attributed to reductions of either food intake or body fat stores, since the former was actually increased and the latter unchanged throughout the study period. Instead, the data are more consistent with a model in which AgRP-neuron activation contributes to cold-induced hyperphagia.

### The increases of energy expenditure and energy intake during cold exposure are rapid in onset

To delineate in greater detail the timing of the onset of feeding and metabolic responses to mild cold exposure, we commenced serial metabolic measurements starting when mice were placed into metabolic cages (to which they had been previously acclimated) that were inside environmental chambers that were either pre-cooled (14°C) or set at room temperature (22°C) in a randomized, cross-over manner (**Figure 2A**). Our findings show that the increase of heat production associated with the change of ambient temperature increased rapidly, such that the thermogenic response needed to maintain core temperature was detected within 5 min (**Figure 2B**) and remained elevated for the duration of the 24-h study (**Figure 2C**). Somewhat surprisingly, we also found that upon acute cold exposure, the increase of food intake required to offset heightened thermogenic demand was equally swift (**Figure 2D**). By comparison, neither respiratory quotient nor ambulatory activity was affected during cold exposure, suggesting that they are unlikely to play an important role in the adaptive response to this challenge (**Supplemental Figure 1**). Like the increase of heat production, the initial increase in the rate of food consumption was maintained throughout both light and dark cycles (**Figure 2E**). Based on the rapidity of this feeding response, we infer that it was unlikely to have resulted from any detectable depletion of body fat or associated circulating signal (e.g., leptin). We interpret these findings as favoring a model in which the perception of cold is sufficient to increase both food intake and energy expenditure rapidly and in parallel, independently of feedback signals that might be subsequently recruited.

**Figure 2.**
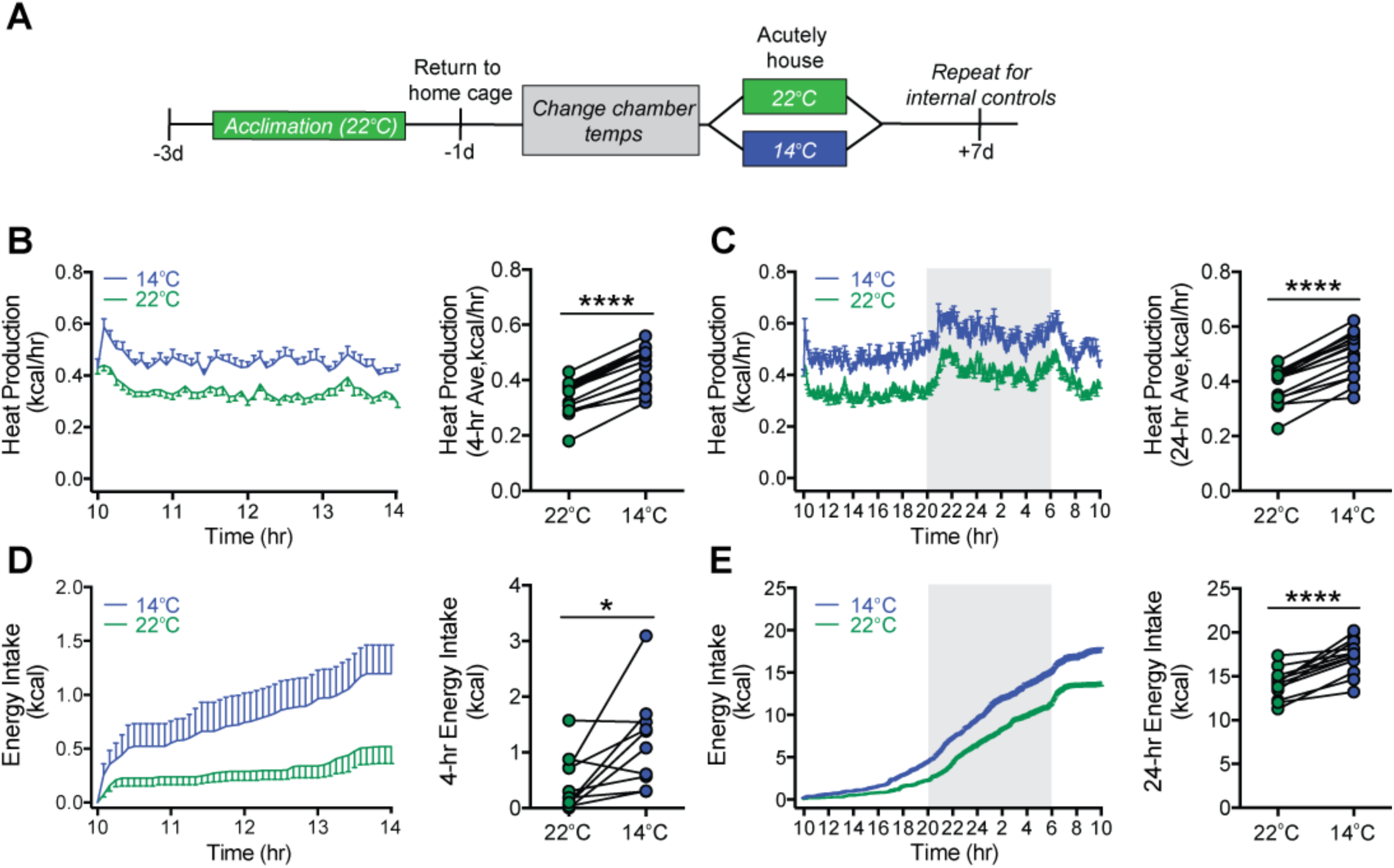
Acute mild cold exposure rapidly increases both energy expenditure and energy intake. (**A**) Adult male wild-type mice were acutely housed in temperature-controlled chambers set to either mild cold (14°C) or, as control, room temperature (22°C). Time-series and mean heat production over (**B**) 4-hr and (**C**) 24-hr, respectively, and time-series and total energy intake over (**D**) 4-hr and (**E**) 24-hr, respectively beginning at 10:00 AM, n=10/group, mean ± SEM. RM-ANOVA and paired Student’s t-test, ****p<0.0001, *p<0.05.

### Mild cold exposure induces Fos in AgRP neurons

We next investigated whether the rapid onset of cold-induced hyperphagia is associated with AgRP neuron activation. To this end, we utilized transgenic AgRP-Cre:GFP mice to enable visualization of Fos protein, a marker of neuronal activation (Kovács, et al., 1998), in AgRP neurons after placement in cages previously set to mild cold (14°C), room temperature (22°C), or thermoneutrality (∼30°C) for 90 minutes with food present. Relative to mice maintained at 22°C, we found that both the total number of Fos+ cells in the ARC (**Figure 3D**) and, specifically, the number of Fos+ AgRP neurons (**Figure 3E-F**) was robustly increased within 90-min of cold-exposure onset. As expected, we also found the expected cold-induced increases in Fos in other thermoregulatory control centers including the parabrachial nucleus (PBN), preoptic area (POA) (Bratincsák & Palkovits, 2004; Geerling, et al., 2016), dorsomedial hypothalamus (DMH) (Hunt, et al., 2009), and rostral raphe pallidus (rRPa) (**Supplemental Figure 2**). Moreover, in contrast to both 14°C and 22°C, both the total number of Fos+ cells in the ARC and the number of Fos+ AgRP neurons was minimal in mice housed at 30°C (**Figure 3D-F**). This Fos response was distributed equivalently across the entire rostral to caudal AgRP neuron population (**Supplemental Figure 3**). Our finding that Fos+ AgRP neuron number increases at temperatures below thermoneutrality (**Figure 3F**) identifies ambient temperature as a potential physiological regulator of AgRP-neuron activity.

**Figure 3.**
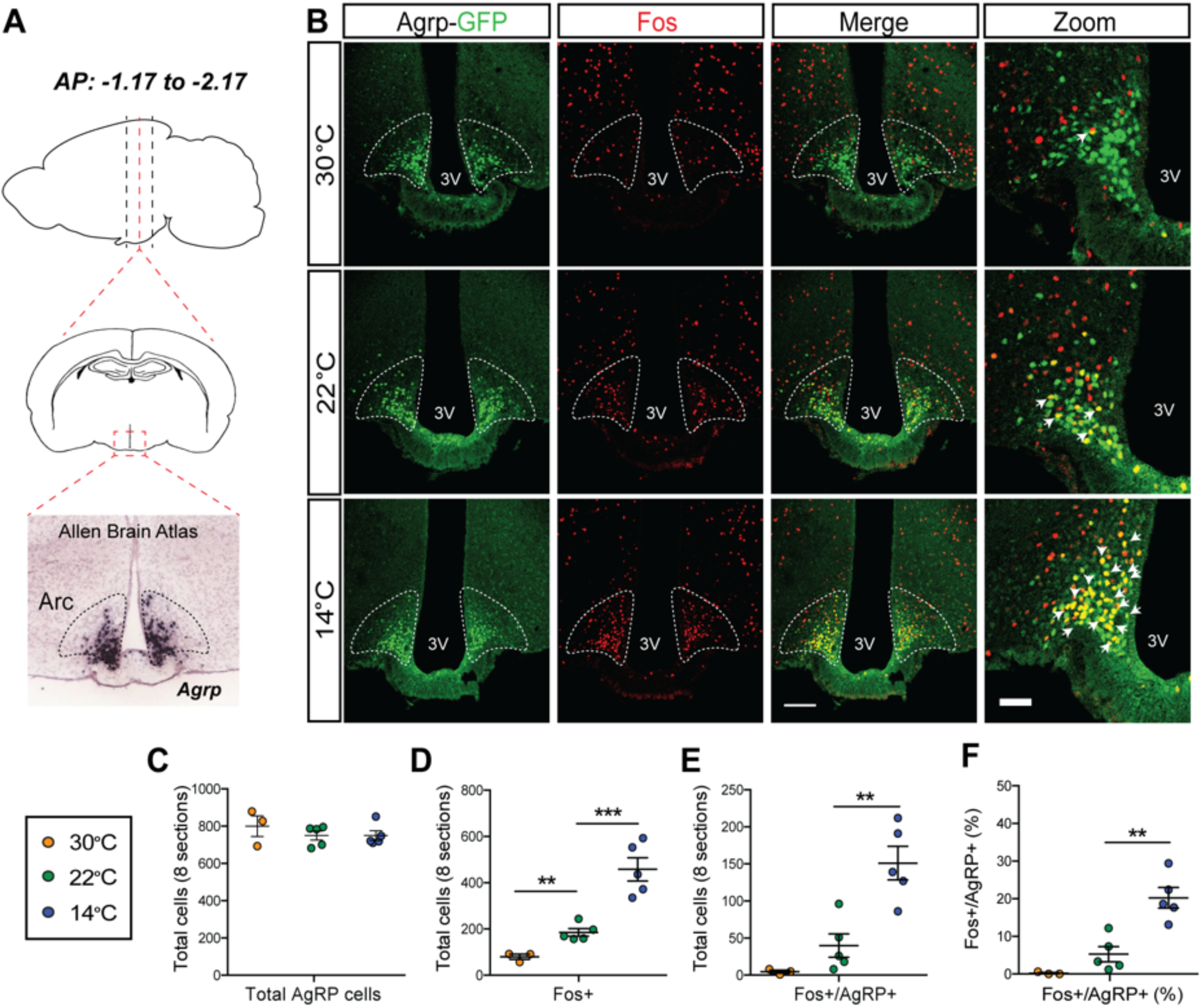
Acute mild cold exposure activates AgRP neurons. (**A**) Representative sagittal and coronal images and *Agrp* hybridization *in situ* from Allen Brain Institute of the ARC. (**B**) Immunohistochemical detection of AgRP-locus driven GFP (green), Fos (red), and colocalization of GFP and Fos (two right panels) in the ARC of AgRP-Cre:GFP mice 90 min after housing at either 14°C or 30°C or 22°C. Quantitation of (**C**) total AgRP^+^ cells, (**D**) total Fos^+^ cells and the total number (**E**) and percent (**F**) of AgRP neurons that co-express Fos across 8 rostral to caudal sections of the ARC (AP: -1.17 to -2.17). Thin bar = 100µm, Thick bar = 50µm, n=3-5/group, mean ± SEM. One-way ANOVA, ***p<0.001, **p<0.01, *p<0.05 vs. 22 °C.

### Mild cold exposure rapidly increases Agrp neuron activity

To definitively establish the timing with which AgRP-neuron activity changes in relationship to cold exposure, we applied fiber photometry of AgRP neurons *in vivo* during a thermal challenge. To this end, AgRP-IRES-Cre mice received a unilateral injection of an adeno-associated virus (AAV) containing a Cre-dependent, genetically-encoded calcium sensor GCaMP6s directed to the ARC (Chen, et al., 2013), followed by implantation of an optical fiber at the injection site (**Figure 4A-B**). After allowing 3 weeks for transgene expression and acclimation, validation of signal quality was performed by examining the AgRP neuron response to food presentation in the fasted state. Consistent with previous observations (Betley, et al., 2015; Chen, et al., 2019; Mandelblat-Cerf, et al., 2015), we found that food presentation strongly and rapidly inhibited AgRP neuron activity, as indicated by a strong reduction in GCaMP signal (**Supplemental Figure 4**). To determine whether exposure to cold affects AgRP neuron activity, mice were placed in a custom-built plexiglass cage modified to enable rapid control over the temperature sensed by the tethered animal (**Figure 4C**). We predicted that if AgRP neurons contribute to cold-induced hyperphagia, exposure to cold would activate AgRP neurons over a time frame that would either precede or coincide with hyperphagia onset (**Figure 2D**). Consistent with this prediction, we found that when the temperature of the platform was rapidly reduced from 30°C to 14°C (within 1 min) an increase in GCaMP activity was detected within seconds, consistent with an increase in AgRP neuronal activity. Moreover, this effect was reversed by raising the temperature from 14°C back to 30°C (**Figure 4D-F**).

**Figure 4.**
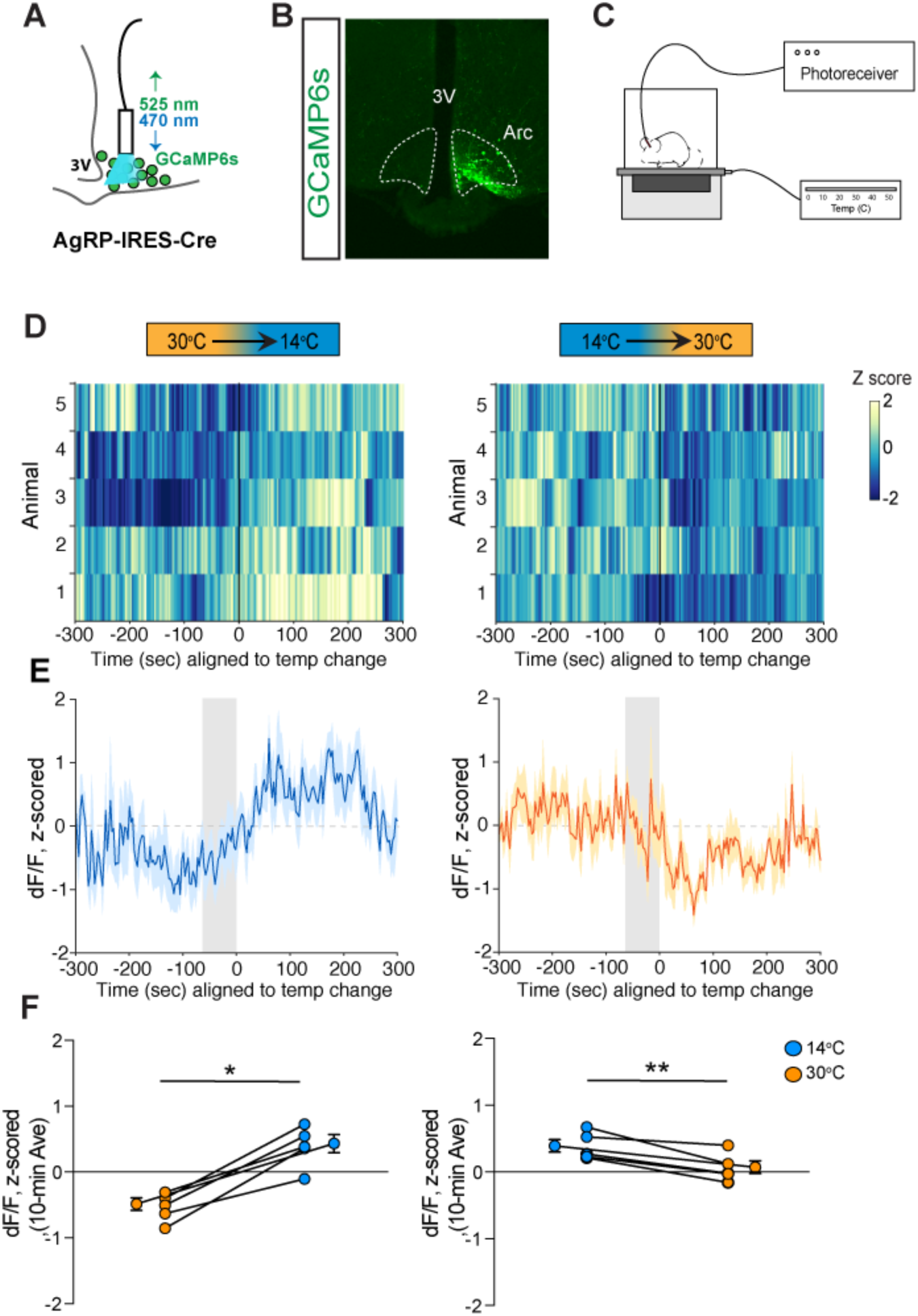
Cold sensation increases AgRP neuron GCaMP activity in a rapidly reversible manner. (**A**) Representative diagram of fiber photometry with fiber placement at ARC. (**B**) Unilateral GCaMP6s expression in AgRP neurons. (**C**) Representative diagram of experimental set up. (**D**) Heat maps of z-scored dF/F, (**E**) trace of averaged z-scored dF/F GCaMP6s signal, and (**F**) quantification of mean differences between 30°C and 14°C GCaMP activity during 10-minute exposure. Gray bar signifies 60-sec temperature-ramp transition, n=5/group, mean ± SEM. Student’s paired t-test, **p<0.01, *p<0.05.

At the end of each study, animals were presented with a food pellet to establish intact inhibition of AgRP neuron GCaMP activity. Although the animals were fed ad libitum prior to the study, a clear inhibition of AgRP neuron GCaMP activity was present comparable to the changes seen during temperature transitions, thus validating the photometry signal (**Supplemental Figure 5**). Taken together, these findings indicate that AgRP neurons are rapidly activated in response to cold exposure.

### AgRP-neuron activity is necessary for cold-induced hyperphagia

Having established that AgRP neurons are rapidly activated in response to mild cold exposure, we next sought to determine whether this response is required for the associated hyperphagia. To test this hypothesis, we utilized a chemogenetic approach in which AgRP-IRES-Cre mice received bilateral microinjections into the ARC of an AAV construct containing a Cre-dependent cassette encoding the inhibitory designer receptor activated by a designer drug (DREADD), hM4Di:eYFP (**Figure 5A**). The transduced neurons can be detected by their eYFP expression and, as expected, it was limited to the ARC (**Figure 5B**). Acclimated animals received an intraperitoneal injection of clozapine-N-oxide (CNO) or its vehicle (saline) in a randomized crossover manner, one hour prior to being placed into metabolic cages housed within temperature-controlled chambers previously set at either 22°C, 30°C or 14°C for measures of food intake and energy expenditure by indirect calorimetry for 4 hours (**Figure 5C**).

**Figure 5.**
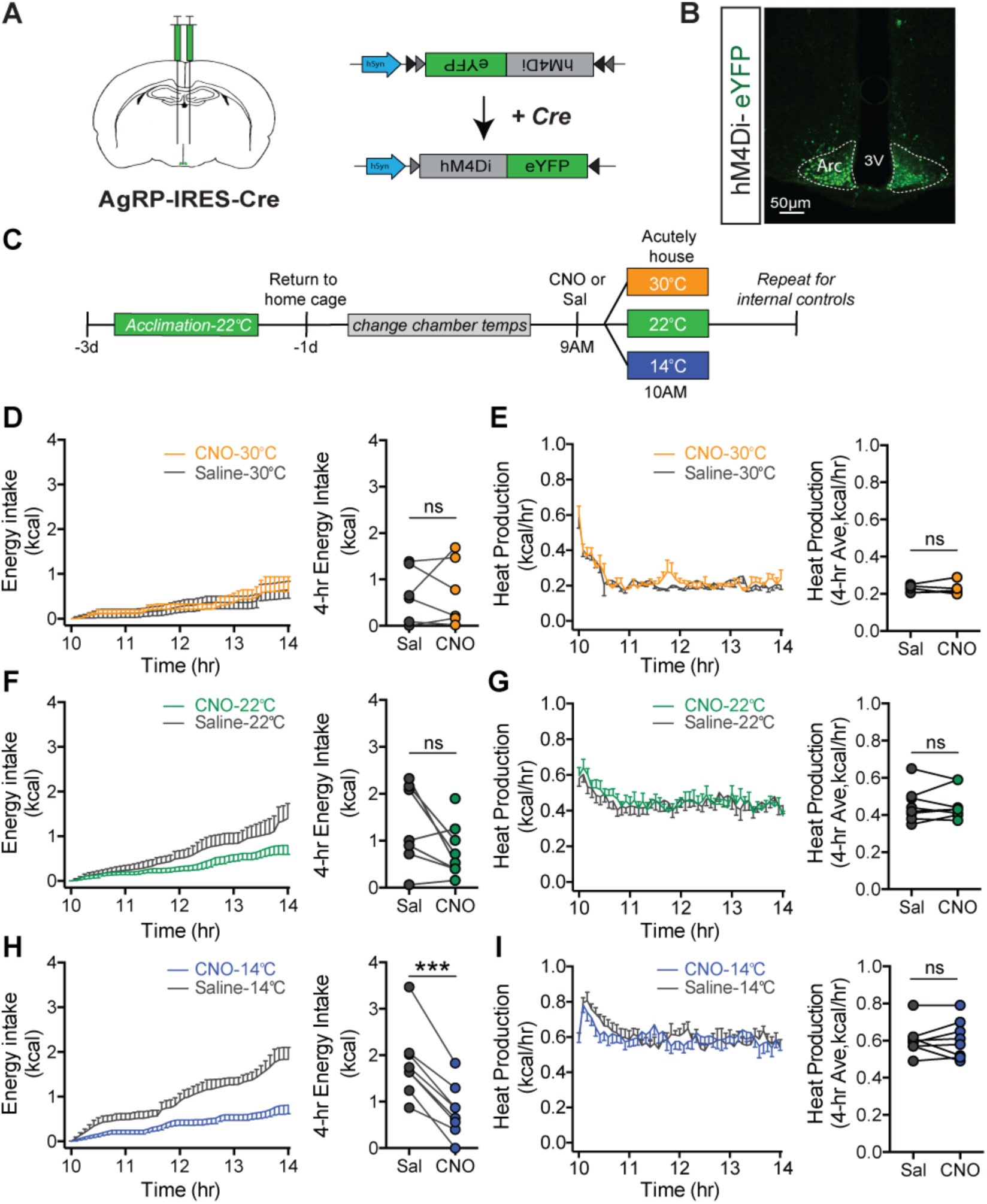
Cold-induced hyperphagia, but not thermogenesis, requires activation of AgRP neurons. (**A**) Schematic depicting strategy for bilateral microinjection of the Cre-dependent inhibitory DREADD (hM4Di) virus into the ARC of AgRP-IRES-Cre mice. (**B**) Detection of bilateral hM4Di-eYFP in transduced AgRP neurons in the ARC. (**C**) Adult male hM4Di-eYFP AgRP-IRES-Cre mice were acclimated to temperature-controlled chambers set to 22°C. Animals were returned to their home cages and chamber temperatures were adjusted overnight. In the morning, animals were dosed i.p. with either CNO or saline one hour prior to being acutely housed at either mild cold (14°C), room temperature (22°C), or thermoneutrality (30°C) in a randomized, crossover manner. (**D, F, H**) 4-hr time-series and mean values for energy intake, and (**E, G, I**) 4-hr time-series and mean values of heat production in hM4Di-eYFP AgRP-IRES-Cre mice that were housed at either 30°C, 22°C or 14°C after receiving an i.p. injection of either saline or CNO n=6-8/group, Mean ± SEM. RM-ANOVA and Student’s t-test, ***p<0.001, *p<0.05 vs. saline.

Based on our Fos and fiber-photometry data (**Figures 3 and 4**), we predicted that the effect of chemogenetic AgRP-neuron inhibition would be robust in mice housed at 14°C, but minimal at 30°C, because AgRP neurons exhibit low activity at thermoneutrality. Consistent with this prediction, we found that whereas inhibition of AgRP neurons had little effect on food intake in mice housed at 30°C (**Figure 5D**), intake was strongly reduced in animals housed at 14°C, while intake was more modestly inhibited by CNO in mice housed at 22°C (**Figure 5F and H**). The latter finding is consistent with room temperature posing a thermal stress to mice (Abreu-Vieira, et al., 2015).

The effect of AgRP-neuron inhibition appears to be selective for energy intake, as the cold-induced increase of heat production was not affected (**Figure 5E, G, I**). In addition, there was no significant effect of AgRP-neuron inhibition on either respiratory quotient or ambulatory activity at any of the three ambient temperatures studied (**Supplemental Figure 6**). Taken together, these findings suggest that AgRP-neuron activation is required for cold-induced hyperphagia, neurocircuits underlying feeding and thermogenic responses to cold are distinct and separable from one another, and some degree of cold stress (including housing at room temperature) is required for food intake to be reduced by chemogenetic, AgRP-neuron inhibition under ad libitum feeding conditions during the light cycle.

## DISCUSSION

The tight coupling that exists between thermoregulation and energy homeostasis (Gordon, 1993; Kaiyala, et al., 2015) enables animals to precisely maintain both core body temperature and body fat mass across a wide range of ambient temperatures, so long as food is available (Kaiyala, et al., 2015; Smith, et al., 1984; Thurlby, et al., 1979). Fundamental to this process is the precise coupling between changes in energy expenditure and energy intake in response to changing requirements for heat production (Armitage, et al., 1984; Kaiyala, et al., 2015; Thurlby, et al., 1979). While cold-induced hyperphagia has been documented across many species, including humans (Johnson & Kark, 1947), the mechanisms underlying this response have received little attention, especially when compared to the considerable literature describing how changes of thermogenesis are coupled to changes of ambient temperature (Madden & Morrison, 2019; Rezai-Zadeh, et al., 2013). This relative lack of research interest may reflect the pervasive view that rather than being an integral component of the thermoregulatory control system, cold-induced hyperphagia is mounted as a compensatory response to the negative-energy balance state induced by cold-induced thermogenesis. Our current data suggest that this perspective should be revised.

Given the well-documented role of AgRP neurons in physiological control of food intake and energy homeostasis (Schwartz, et al., 2000; Timper, et al., 2017; Varela, et al., 2012), we hypothesized that cold-induced hyperphagia is driven at least in part by activation of these neurons. We report that in normal mice, AgRP neurons are rapidly activated by cold exposure and that intact cold-induced hyperphagia requires this response, since it was blocked by acute chemogenetic AgRP neuron silencing. We also show that the magnitude of food-intake reduction induced by chemogenetic silencing of these neurons varies inversely with increasing ambient temperature, such that the effect is more pronounced in colder environments relative to thermoneutral conditions in which AgRP-neuronal activity is lower. Taken together, we conclude that under our experimental conditions (*e.g.*, in the absence of negative energy balance induced, for example, by caloric restriction), the adaptive increase of food intake during acute cold exposure depends upon increases in AgRP-neuron activity.

AgRP neurons are among the most studied hypothalamic neuronal populations involved in energy homeostasis. These GABAergic neurons co-express the potent orexigenic neuropeptide Y (NPY) and are found exclusively in the ARC (Hahn, et al., 1998). An adjacent neuronal population expresses pro-opiomelanocortin (POMC), and whereas AgRP-neuron activation stimulates feeding, POMC-neuron activation has the opposite effect (Schwartz, et al., 2000; Xu, et al., 2011). Furthermore, AgRP and POMC neurons are reciprocally regulated by leptin (Schwartz, et al., 2000; Sohn, et al., 2013) and other humoral signals, such that the effect of energy restriction to reduce body fat stores (and plasma leptin levels) causes both activation of AgRP neurons and inhibition of POMC neurons, a combined response that drives feeding until body fat stores and plasma leptin levels return to baseline values (Schwartz, et al., 2017). These considerations raise the possibility that the feeding response to cold is mounted as a compensatory response triggered by increased thermogenesis, which in turn causes depletion of body fuel stores and associated reduction of adiposity negative feedback (e.g., falling plasma leptin levels) (Bing, et al., 1998; Puerta, et al., 2002; Ricci, et al., 2000; Tang, et al., 2009). However, the rapidity with which AgRP neurons are activated in response to cold exposure seems inconsistent with this type of negative-feedback control. Nevertheless, humoral negative-feedback signals could certainly be recruited over time and play a role in sustained feeding responses to a cold challenge, even if they are not relevant to the initial hyperphagic response. Thus, investigation of the role of POMC neurons or other neuronal cell types involved in homeostatic control of feeding is warranted.

Recent work has established that in addition to negative-feedback regulation by humoral signals relevant to body fuel stores, AgRP-neuron activity is also influenced by “feed-forward” signals that promote homeostasis by anticipating future need (Chen, et al., 2016; Lowell, 2019). Among relevant feed-forward signals are cues from the environment that anticipate eating (e.g., the sight or smell of food), which are presumably communicated indirectly to AgRP neurons via cortical areas that process the relevant sensory input (Livneh, et al., 2017). How this information is ultimately transmitted to AgRP neurons is unknown, but synaptic relays in the hypothalamic paraventricular nucleus (PVN) (Krashes, et al., 2014), dorsomedial nucleus (DMH) (Garfield, et al., 2016), and POA may play a role. The key point is that although food intake driven by AgRP-neuron activation is commonly associated with states of negative energy balance and/or falling leptin levels, these neurons can also be activated by sensory-related stimuli (Chen, et al., 2016; Garfield, et al., 2016; Krashes, et al., 2014).

These considerations have relevance to our finding that AgRP neurons are rapidly activated following exposure to a cold environment, and that the activity of these neurons at thermoneutrality is lower. These observations are consistent with previous findings in neonates suggesting that AgRP neurons are regulated by ambient temperature. Specifically, exposure of P10 mice to a warm environment rapidly suppresses AgRP-neuron activity, whereas pup isolation (with reduced thermal insulation from dam and littermates) increases AgRP-neuron activity (Zimmer, et al., 2019). Moreover, the time course of increased AgRP-neuron activity coincides with the increase of both energy expenditure and energy intake elicited by cold exposure. By commencing continuous monitoring upon placing mice in a pre-cooled cage (rather than waiting for the cage to gradually cool to the desired temperature), we were able to show that energy expenditure increases rapidly in response to cold exposure – such that the thermogenic response needed to maintain core body temperature for the duration the study was achieved within 5 min. Even more impressive is that cold-induced hyperphagia has a similarly rapid onset, at least in mice studied during the mid-light cycle, when they are normally inactive and consume only a small percentage of their daily calories (Ellacott, et al., 2010). We interpret the rapidity and synchrony with which exposure to a cold environment activates AgRP neurons and engages feeding responses as being consistent with control via a “feed-forward” mechanism, rather than control by negative-feedback control, and future studies are warranted to test this hypothesis.

Our finding that blockade of cold-induced hyperphagia by chemogenetic inhibition of AgRP neurons occurred despite having no effect on energy expenditure suggests that cold-induced AgRP-neuron activation is required for the feeding, but not thermogenic response to this challenge. We also report that AgRP-neuron inhibition is without effect on either respiratory quotient or ambulatory activity in cold-exposed mice. Based on these findings, we conclude that the primary contribution made by AgRP-neuron activation to the adaptive response to cold exposure is to drive hyperphagic feeding.

In conclusion, we report that AgRP-neuron activation occurs rapidly during cold-exposure, and that this response plays a key role to drive the associated hyperphagic, but not the thermogenic, response to this stimulus. Further, we demonstrate that the contribution made by AgRP-neuron activation to food consumption is dependent on ambient temperature, constituting a large fraction of intake during cold exposure while playing a lesser role in a thermoneutral environment. In addition to advancing our understanding of the thermoregulatory system, insights into the neurocircuitry linking thermoregulation to AgRP-neuron activity may help to identify novel strategies for obesity treatment by blunting the associated hyperphagic response.

## MATERIALS AND METHODS

### Animals

All procedures were performed in accordance with NIH Guide for the Care and Use of Laboratory Animals and were approved by the Institutional Animal Care and Use Committee at the University of Washington. Mice were individually housed in a temperature-controlled room with either a 12:12 hr or 14:10 hr light:dark cycle under specific-pathogen free conditions and were provided with *ad libitum* access to water and fed a standard laboratory chow (5001; 13% kcal fat, LabDiet, St. Louis, MO), unless otherwise stated. Adult male C57Bl/6 wild-type mice were obtained from Jackson Laboratories, ME, while AgRP-IRES-Cre mice were kindly provided to us by Dr. Streamson Chua, Jr (Albert Einstein College of Medicine) and have been previously described (53). The AgRP-Cre:GFP knock-in mice (version 2, v2) used in this study were generated by Dr. Richard Palmiter (University of Washington) by replacing the Cre:GFP cassette of the original line (Sanz et al., 2015) with a new cassette designed to have attenuated expression of Cre:GFP. The new cassette differs from the original by: (a) removing the nuclear localization signal from Cre, (b) using a non-optimal initiation codon, (c) removing part of the N-terminal sequence of Cre, and (d) adding a 3’ untranslated region from the Myc gene that promotes a short mRNA half-life. Crossing mice with this new (v2) line with a conditional reporter line of mice (Gt(ROSA)26-LSL-TdTomato, Allen Institute Seattle WA) gives faithful expression only in AgRP neurons, unlike the original line (Sanz, et al., 2015) which occasionally resulted in ectopic expression in many parts of the brain. The new targeting construct was electroporated into G4 ES cells (C57Bl/6 x 129/SV) and clones with correct targeting were identified by Southern blot of DNA digested with BamH1; 6 of 48 clones analyzed were correctly targeted. Three of these clones were injected into blastocysts from C57Bl/6 mice and one of them gave chimeras with a high percentage of agouti color. Those chimeras were bred with Gt(ROSA)26-FLPer mice to remove the frt-flanked Neo gene. Thereafter, the mice were backcrossed to C57Bl/6 mice for more than 6 generations before our experiments were performed.

### Viral constructs

Chemogenetic inhibition of AgRP neurons was achieved by microinjection of an AAV containing Cre-dependent cassette for the inhibitory DREADD, AAV1-CAG-DIO-hM4Di-YFP-WPRE-bGHpA (hM4Di: kindly provided by the laboratory of Dr. Larry Zweifel, University of Washington), into brain areas containing Cre-expressing neurons (e.g., the ARC of AgRP-IRES-Cre mice). Activation of the DREADD receptor was induced by intraperitoneal administration of the agonist, clozapine-N-oxide (CNO, 1 mg/kg, ip). For fiber-photometry experiments, we utilized an AAV containing a Cre-dependent cassette for the genetically-encoded calcium indicator, GCaMP6s, AAVDJ-EF1a-DIO-GCaMP6s-WPRE (UNC Viral Core, Chapel Hill, NC).

### Surgery

Stereotaxic viral injections were performed as described (Faber et al., 2018; Meek et al., 2016). Briefly, animals were anesthetized using 1-3% isoflurane, their head shaved and placed in three-dimensional stereotaxic frame (Kopf 1900, Cartesian Research Inc., CA). For inhibitory DREADD experiments, the skull was exposed with a small incision, and two small holes were drilled for bilateral microinjection (400 nl/side) of the inhibitory DREADD (hM4Di) AAV into the ARC of AgRP-IRES-Cre mice at stereotaxic coordinates based on the Mouse Brain Atlas: A/P: -1.2, M/L: +/-0.3, D/V: -5.85 (Franklin & Paxinos, 2013). For fiber photometry experiments, AAV was injected using a unilateral and angled approach (A/P: -1.8, D/V: -5.85, 12° angle from midline). After viral injections, a fiberoptic ferrule (0.48 NA, Ø400 *μ*m core; Doric Lenses, Quebec, Canada) was implanted using the same coordinates. All microinjections were performed using a Hamilton syringe with a 33-gauge needle at a flow rate of 100 nL/min (Micro4 controller, World Precision Instruments, Sarasota, FL), followed by a 5-min pause and slow withdrawal. Animals received a peri-operative dose of buprenorphine hydrochloride (0.05 mg/kg sc; Reckitt Benckiser, Richmond, VA). After surgery, mice were allowed 3 weeks to recover to maximize virally-transduced gene expression and to acclimate animals to handling and experimental paradigms prior to study. Expression and fiber placement were verified post hoc in all animals, and any data from animals in which the transgene expression and/or fiber was located outside the targeted area were excluded from analysis. Five out of 10 mice were excluded due to improper targeting of fiber or viral injection.

### Body composition analysis

Determination of body fat and lean mass was performed using quantitative magnetic resonance spectroscopy (EchoMRI™, Houston, TX) in conscious mice using the NIDDK-funded Nutrition Obesity Research Center Energy Balance Core (Taicher, et al., 2003).

### Indirect calorimetry

For indirect calorimetry studies, C57Bl/6 mice were acclimated to metabolic cages after which energy expenditure was measured using a computer-controlled indirect calorimetry system (Promethion®, Sable Systems, Las Vegas, NV) as described (Kaiyala et al., 2015; 2012; Kaiyala, et al., 2016). For each animal, O_2_ consumption and CO_2_ production were measured for 1 min at 5-min (acute studies) or 10-min (chronic studies) intervals. Respiratory quotient (RQ) was calculated as the ratio of CO_2_ production to O_2_ consumption. Energy expenditure was calculated from VO_2_ and VCO_2_ data using the Weir equation (Weir, et al., 1949). Ambulatory activity was measured continuously with consecutive adjacent infrared beam breaks in the x-, y- and z-axes were scored as an activity count that was recorded every 5 or 10 min. Data acquisition and instrument control were coordinated by MetaScreen v.1.6.2 and raw data was processed using ExpeData v.1.4.3 (Sable Systems) using an analysis script documenting all aspects of data transformation.

### Core body temperature monitoring

Adult male C57Bl/6 mice received body temperature transponders implanted into the peritoneal cavity (Starr Life Science Corp, Oakmont, PA) and were allowed a one-week recovery period. Animals were then acclimated to metabolic cages enclosed in temperature- and humidity-controlled cabinets (Caron Products and Services, Marietta, OH) prior to study. Signals emitted by body-temperature transponders were sensed by a platform receiver positioned underneath the cage and analyzed using VitalView software as described (Kaiyala, et al., 2015; Kaiyala, et al., 2016).

### Immunohistochemistry

For immunohistochemical studies, animals were overdosed with ketamine:xylazine and perfused with 1X phosphate-buffered saline (PBS) followed by 4% (v/v) paraformaldehyde in 0.1M PBS. Brains were removed and post-fixed for 4 hr in paraformaldehyde followed by sucrose (30%) dehydration and embedding in OCT blocks. Free-floating coronal sections were obtained via Cryostat at 35-μm thickness and stored in 1X PBS with 0.02% sodium azide at 4°C. Free-floating sections were then washed at room temperature in phosphate-buffered saline with 0.1% Tween 20 or 0.4% Triton-X 100 (PBS-T) for one hour, followed by a blocking buffer (5% normal donkey serum, 1% bovine serum albumin in 0.1M PBS-T with 0.01% sodium azide) for an additional hour with rocking. Sections were then incubated 24-48 hr at 4°C with polyclonal rabbit anti-cFos (Ab-5 (4-17), 1:10,000, RRID:AB_2106755; Millipore, Burlington, MA) and/or Chicken anti-GFP (Abcam, ab13970, 1:10,000, RRID:AB_300798; Cambridge, UK) in blocking buffer, followed by PBS-T washes at room temperature. Sections were then incubated in secondary donkey anti-rabbit Alexa 594 or donkey anti-chicken Alexa 488 (1:500, Jackson ImmunoResearch Laboratories, West Grove, PA) in blocking buffer overnight at 4°C, followed by PBS-T washes. Sections were stained with DAPI (1:10,000, Sigma, St. Louis, MO) for 30 minutes, followed by a final set of washes and mounting with Fluoromount-G (Thermo Fisher Scientific, Wilmington, DE) or prepared polyvinyl acetate (PVA).

### qRT-PCR

To quantify specific hypothalamic mRNA transcripts, mice were sacrificed at study completion and the hypothalamus rapidly dissected and flash frozen. Individual tissue samples were dounce-homogenized and RNA was isolated using Qiagen RNeasy Micro Kit (Kit# 74004, Hilden, Germany) and isolated RNA concentrations were quantified by Nanodrop (Thermo Fisher Scientific, Wilmington, DE). qRT-PCR was performed using SYBR Green One-Step (Kit# 600825, Agilent, Santa Clara, CA). qRT-PCR data were analyzed using the Sequence Detection System software (SDS Version 2.2; Applied Biosystems, Foster City, CA). Expression levels of each gene were normalized to a housekeeping gene (18S RNA) and standard curve. Non-template controls were incorporated into each PCR run. Oligonucleotides selected were: Agouti-related peptide (*Agrp*): For: 5’-ATGCTGACTCGAATGTTGCTG-3’, Rev: 5’-CAGACTTAGACCTGGGAACTCT-3’, Pro-opiomelanocortin (*Pomc*): For: 5’-CAGTGCCAGGACCTCAC-3’, Rev: 5’-CAGCGAGAGGTCGAGTTTG-3’, Neuropeptide Y (*Npy*): For: 5’-CTCCGCTCTGCGACACTAC-3’, Rev: 5’-AGGGTCTTCAAGCCTTGTTCT-3’, Pro-melanin concentrating hormone (*Pmch*): For: 5’-GAATTTGGAAGATGACATAGTAT-3’, Rev: 5’-CCTGAGCATGTCAAAATCTCTCC-3’, 18S Ribosomal RNA (*18S*): For: 5’-CGGACAGGATTGACAGATTG-3’, Rev: 5’-CAAATCGCTCCACCAACTAA-3’.

### Fiber photometry

Mice expressing GCaMP6s in AgRP neurons were connected to a fiber-photometry system to enable fluorometric analysis of real-time neuronal activity. Briefly, for calcium recording *in vivo*, two excitation wavelengths (470 nm and 405 nm isosbestic) were used to indicate calcium-dependent and calcium-independent (i.e., due to bleaching and motion artifacts) GCaMP6s fluorescence, respectively. Light was delivered via fiber-coupled LEDs (LED lights: M470F3 and M405FP1, LED driver: DC4104, Thorlabs, Newton, NJ) and modulated by a real-time amplifier (RZ5P, Tucker-Davis Technology (TDT), Alachua, FL) at non-divisible frequencies (331 Hz and 231 Hz, respectively) to prevent signal interference between the channels. Excitation lights were bandpass filtered (475 ± 15nm, 405 ± 5nm; iFMC4, Doric Lenses, Quebec, QC, Canada) and the combined excitation light delivered through a fiberoptic patch cord (M75L01,Thorlabs) connected to a rotary joint (FRJ, 0.48 NA, Ø400*μ*m core; Doric Lenses) to prevent fiberoptic torsion during animal movement. A final connector patch cord (MFP, 0.48 NA, Ø400 *μ*m core; Doric Lenses) was connected to the implanted fiberoptic via a ceramic mating sleeve (ADAL1, Thorlabs). Emitted light was collected through the same patch cord, bandpass filtered (525 ± 25 nm; iFMC4) and transduced to digital signals by an integrated photodetector head. Electrical signals were sampled at a rate of 1017.25 Hz and demodulated by the RZ5P real-time processor. Experiments were controlled by Synapse software (TDT).

Custom MATLAB scripts were developed for analyzing fiber photometry data. The isosbestic 405-nm excitation control signal was subtracted from the 470-nm excitation signal to remove movement artifacts from intracellular calcium dependent GCaMP6s fluorescence. Baseline drift was evident in the signal due to slow photobleaching artifacts, particularly during the first several minutes of each recording session. A double exponential curve was fit to the raw trace of temperature-ramping experiments while a linear fit was applied to the raw trace of food presentation experiments and subtracted to correct for baseline drift. After baseline correction, dF/F was calculated as individual fluorescence intensity measurements relative to median fluorescence of entire session for 470nm channel and z-score normalized. Z-score normalized dF/F for each temperature (30 or 14°C) were limited to the 10-min period either before or after the 1-min ramp either between the 14°C to 30°C transition or the 30°C to 14°C transition.

### Measuring determinants of energy balance

To examine the effect of chronic mild, cold exposure on comprehensive measures of energy homeostasis, mice were acclimated to metabolic cages housed within temperature- and humidity-controlled chambers, and either remained at 22°C or they were exposed to 14°C for 5 days. During this period, continuous measures of energy expenditure, respiratory quotient, ambulatory activity and energy and water intake were recorded, and body composition was determined both before and after the thermogenic challenge.

To examine the effect of acute changes in ambient temperature on energy intake, wild-type mice (mean BW: 24.59 ± 0.31 g) were acclimated to metabolic cages housed within temperature- and humidity-controlled chambers. One-hour fasted mice were placed directly into metabolic cages housed within environmental chambers that were pre-set overnight at either 22°C or 14°C at 10 am for 24 hr for continuous measures of energy expenditure, respiratory quotient, ambulatory activity and energy intake. Animals were studied using a randomized cross-over design in which housing conditions were separated by at least 72 hr, with each animal serving as its own control.

### Measuring Fos induction

To determine whether acute cold exposure induces Fos in AgRP neurons, sated AgRP-Cre:GFP mice were housed in temperature-controlled chambers pre-set overnight at either 22°C, 30°C, or 14°C for 90 min. Although food is typically removed during the period prior to Fos quantitation in this type of study (so as to minimize the impact that variation in food intake can have on the activity of AgRP neurons), we opted not to do so here, based on the concern that cold-exposed mice would experience a state of negative energy balance greater than occurred in controls housed at room temperature (owing to a comparatively greater rate of energy expenditure), and that this might confound data interpretation. Animals were then anesthetized and perfused as described below. Immunostaining for Fos was quantified by imaging at either 10X or 20X on a Leica SP8X confocal system with support from the University of Washington, W.M. Keck Microscopy Center. Images were merged using ImageJ (Fiji, NIH) and threshold adjusted to minimize nonspecific background fluorescence. Fos+ cells were then identified and counted in 35-µm sections obtained serially across the full rostral to caudal axis of the arcuate nucleus using the ‘analyze particles’ feature, such that consistent fluorescence and size thresholds were used throughout, as previously described (Faber, et al., 2018). For measures of Fos induction in additional thermoregulatory nuclei, 2-3 sections from each region were counted for each subject.

### Measuring AgRP-neuron activity

To assess the impact of cold exposure on AgRP-neuronal activity, mice were acutely housed on a custom thermal platform. Baseline GCaMP6s fluorescence signals was set to similar levels across animals by adjusting the intensities of the 470-nm and 405-nm LEDs, and baseline recording was measured for 5 min. For temperature-challenge studies, mice were placed in a small custom-built plexiglass temperature chamber (3” x 6” x 6”) that was constructed to enclose a Peltier cooler platform that could be controlled by an external controller (TE Tech, TC720) as described (Tan, et al., 2016). Animals were acclimated on 3 separate days to tether and Peltier platform. For temperature ramp studies, animals were attached to tether and allowed 2-3 min for photobleaching before photometry recording is initiated. Temperature-ramp studies were designed to test GCaMP6s activity in AgRP neurons when transitioning from 30°C to 14°C or 14°C to 30°C. Animals were held at 30°C for 10 min before transitioning to 14°C for 10 min before returning to 30°C with each temperature ramp repeated twice during a recording session. Transitions between temperatures were all set to 60 s. Experiments were aligned to initial 22°C. As an added positive control, at the end of each study, animals were presented with a food pellet, which, led to an expected reduction in AgRP-neuron activity.

Prior to study, mice were acclimated to experimental procedures and AgRP-calcium responses to food presentation in a fast-refeed paradigm was used as a positive control to indicate successfully targeted animals. Briefly, overnight-fasted animals that failed to demonstrate ≥10% reduction in ΔF/F in response to chow presentation were identified as surgical misses and excluded from further study; post hoc IHC validated poor viral expression and/or fiber-optic placement in these animals.

### Cold-induced activation of AgRP neurons

To determine whether activation of AgRP neurons is required for cold-induced hyperphagia, AgRP-IRES-Cre mice (mean BW: 36.10 ±1.06 g) received a bilateral microinjection of the inhibitory DREADD (hM4Di) directed to the ARC. After 3 weeks for recovery from surgery, acclimation to injection/handling, and expression of the transgene, animals were acclimated to environmental chambers to minimize stress. On the day prior to the experiment, animals were returned to their previous home cage and the environmental chambers were allowed 16 hr to come to temperature (30°C, 22°C, or at 14°C). On the experimental day, food was removed at 9 AM and mice received a pre-treatment injection of either CNO (1 mg/kg, ip) or saline in a randomized cross-over manner. At 10AM, mice were placed into metabolic cages housed within environmental chambers set at either 30°C, 22°C, or 14°C with continuous measures of energy intake and energy expenditure recorded over a 4-hr period.

### Statistics

Results are expressed as mean ± SEM. Significance was established at p < 0.05, two-tailed. For statistical comparisons involving core temperature, energy expenditure, ambulatory activity, respiratory quotient or energy intake, data obtained during chronic cold exposure (14°C) relative to room temperature (22°C) were reduced into average light and dark photoperiods for each mouse. A group by ambient temperature ANOVA with least significance difference pairwise tests compared mean values between groups. Where applicable, time-series data were analyzed using treatment by time-mixed factorial ANOVA or linear-mixed model analysis controlling for within subjects correlated data using a random intercept model with treatment and time as fixed effects (Fitzmaurice, et al., 2012). All post hoc comparisons following one-way ANOVA were determined using Sidak correction for multiple comparisons. A two-sample unpaired Student’s t-test was used for two-group comparisons and a paired t-test for within group comparisons. Statistical analyses were performed using Statistica (version 7.1; Statsoft, Incl, Tulsa, OK), SPSS (SPSS version 23, IBM Corp., Somers, NY), R (v 3.6.2, R Core Team 2016), GraphPad Prism (version 7.0; La Jolla, CA), and MATLAB (Mathworks; Natick, MA) (https://github.com/christianepedersen/Cold_Hyperphagia_Requires_AGRP).

## Abbreviations

AgRP: Agouti-related peptide;
ARC: arcuate nucleus;
NPY: neuropeptide Y;
POMC: pro-opiomelanocortin;
PMCH: pro-melanin concentrating hormone;
MPA: medial preoptic area;
(VMPO): ventromedial preoptic area;
DMH: dorsomedial hypothalamus;
PVN: paraventricular nucleus;
PBN: parabrachial nucleus;
rRPa: rostral raphe pallidus;
AAV: adeno-associated virus
GCaMP6s: genetically-encoded calcium indicator, version 6s;
DREADD: designer receptor activated by a designer drug;
hM4Di: inhibitory DREADD receptor;
CNO: clozapine-N-oxide (CNO).

## Acknowledgments

The authors gratefully acknowledge Dr. Larry Zweifel (University of Washington) for providing the inhibitory DREADD virus. This work was supported by National Institutes of Health Grants DK089056 (to G.J.M.), DK083042 and DK101997 (to M.W.S.), R37DA033396 (to M.R.B.), R01-DA-24908 (to R.D.P), the NAPE Center Imaging and Circuits Core (P30 DA048736), the University of Washington W.M. Keck Microscopy Center (S10 OD016240), the Nutrition Obesity Research Center (DK035816), the Diabetes Research Center (DK17047), F31 DK113673 and T32 GM095421 (to C.L.F.), and the Nutrition, Obesity and Atherosclerosis Training Grant (T32 HL007028) at the University of Washington (to J.D.D.), a Dick and Julia McAbee Endowed Fellowship (to J.D.D.) and a American Diabetes Association Innovative Basic Science Award (ADA 1-19-IBS-192, to G.J.M.) and Fellowship Grant (ADA 1-19-PDF-103, to J.D.D.).

## Author Contributions

Conceptualization was performed by J.D.D., J.M.S., M.W.S and G.J.M. Data curation was performed by J.D.D., B.P., J.T.N., M.A.T., and G.J.M. Funding acquisition was performed by J.D.D, J.M.S., M.W.S., and G.J.M. Formal analysis was performed by J.D.D., C.P., K.J.K. and G.J.M. Methodology was performed by J.D.D., C.L.F., C.P., S.A.L., K.O., R.D.P., and G.J.M. Project administration was performed by J.D.D. and G.J.M. Validation was performed by J.D.D., B.P., and V.D. Writing – review and editing – was performed by J.D.D., C.L.F., J.M.S., R.D.P., M.R.B., M.W.S., and G.J.M.

## SUPPLEMENTAL FIGURES

**Supplemental Figure 1 (Related to Figure 2).**
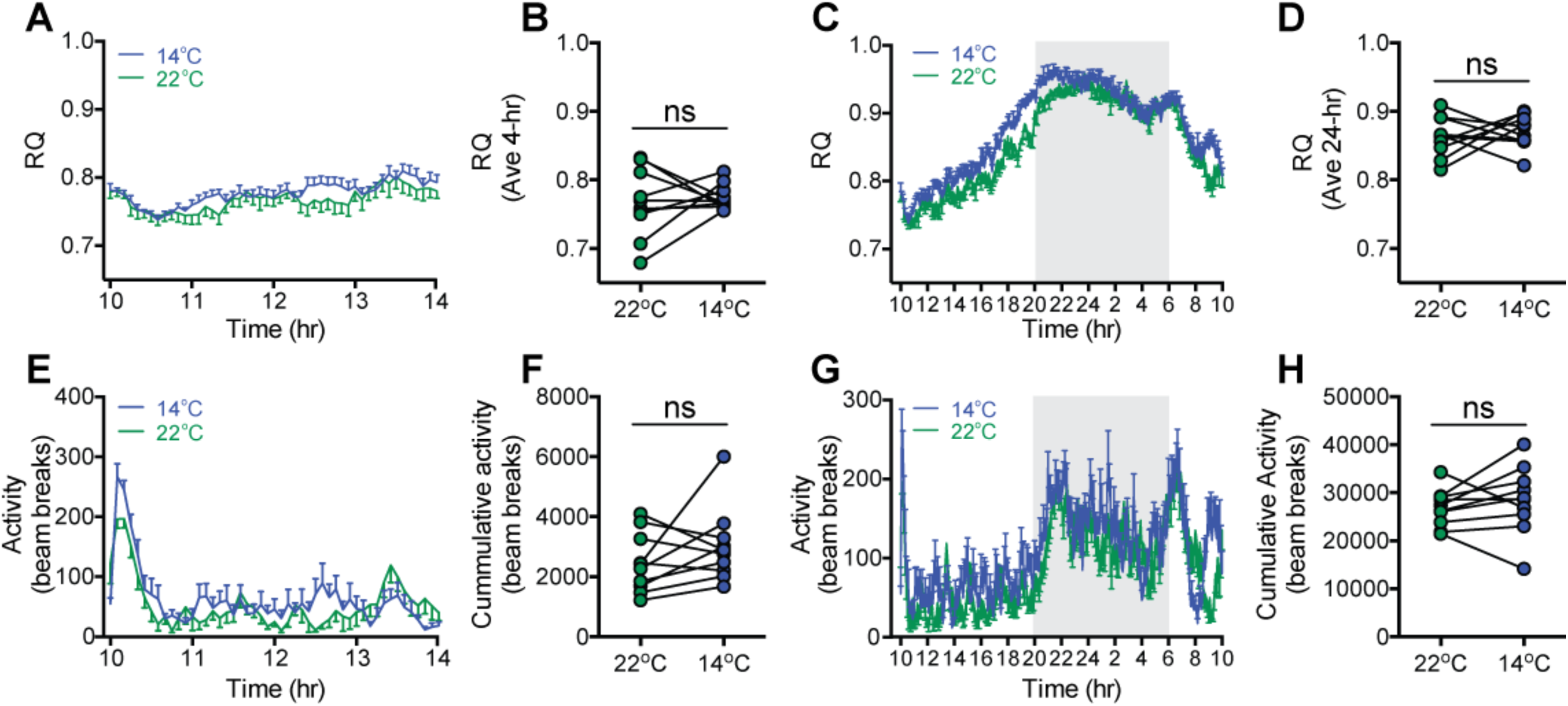
Effect of acute mild cold exposure on respiratory quotient and ambulatory activity. (**A, C**) Time-series and (**B, D**) mean respiratory quotient (RQ) over 4-hr and 24-hr, respectively, and (**E, G**) time-series and (**F, H**) total ambulatory activity over 4-hr and 24-hr, respectively, in adult male wild-type mice moved into housing at either 14°C or 22°C beginning at 10:00 AM, n=10/group, mean ± SEM.

**Supplemental Figure 2 (Related to Figure 3).**
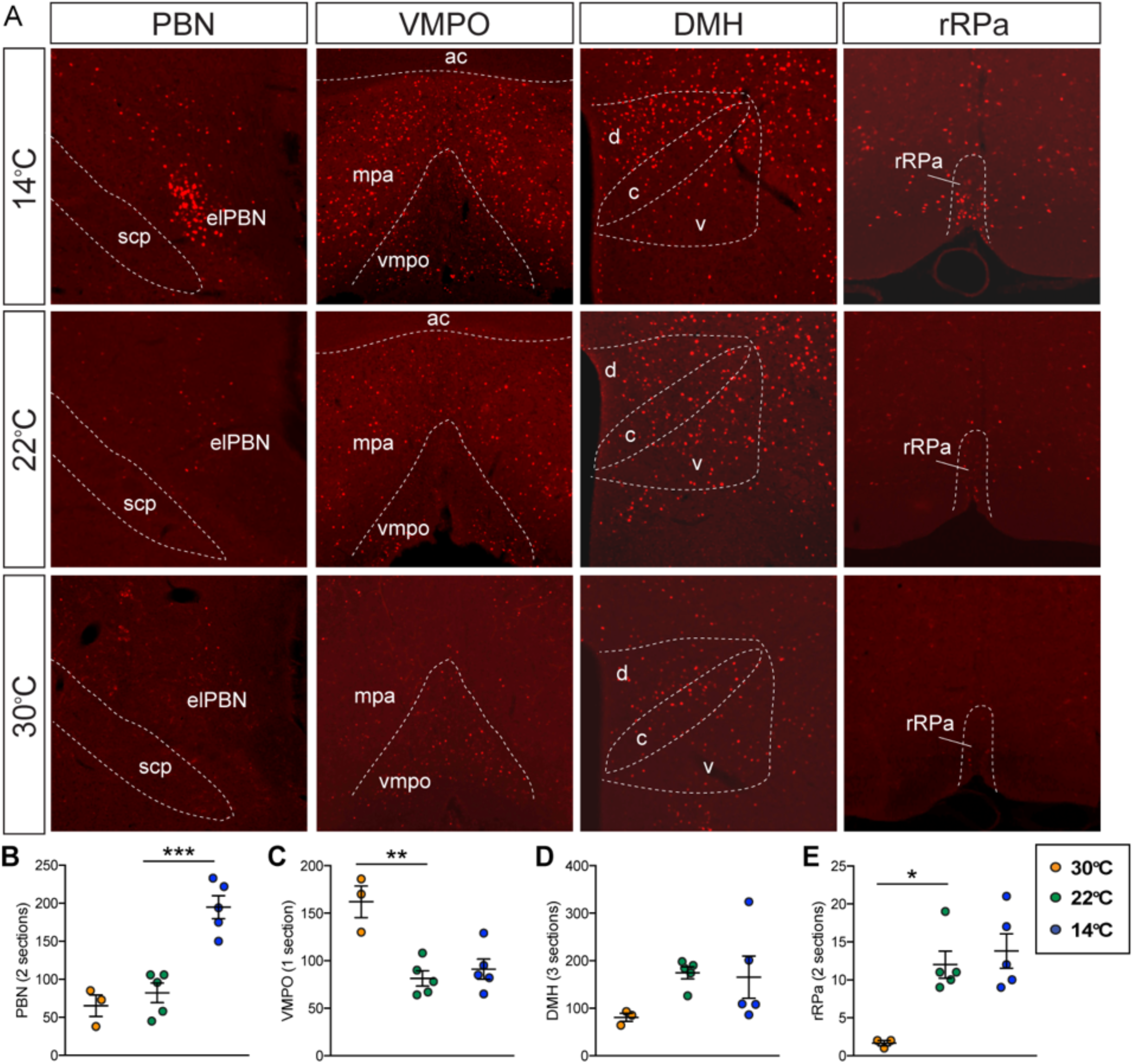
Fos induction in known thermoregulatory brain regions. (**A**) Representative images of Fos induction in known thermoregulatory brain regions. Quantification of Fos induction in (**B**) parabrachial nucleus (PBN), (**C**) ventromedial preoptic area (VMPO) and medial preoptic area (MPA), (**D**) dorsomedial hypothalamus (DMH) and (**E**) rostral raphe pallidus (rRPa) following 90-min of exposure to either 30°C, or 22°C or 14°C, n=3-5/group. Mean ± SEM, One-way ANOVA, ***p<0.001, **p<0.01, *p<0.05 vs. 22°C.

**Supplemental Figure 3 (Related to Figure 3).**
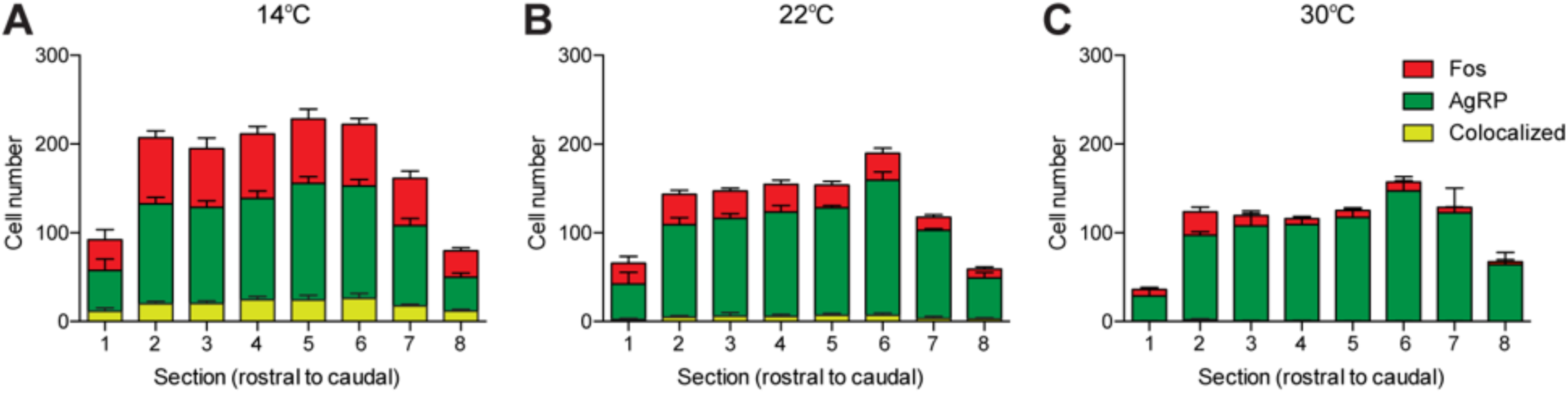
Fos induction in AgRP neurons across entire ARC. Fos induction in AgRP neurons across entire rostral to caudal axis of ARC after exposure to 90-min at (**A**) 14°C, (**B**) 22°C, or (**C**) 30°C.

**Supplemental Figure 4 (Related to Figure 4).**
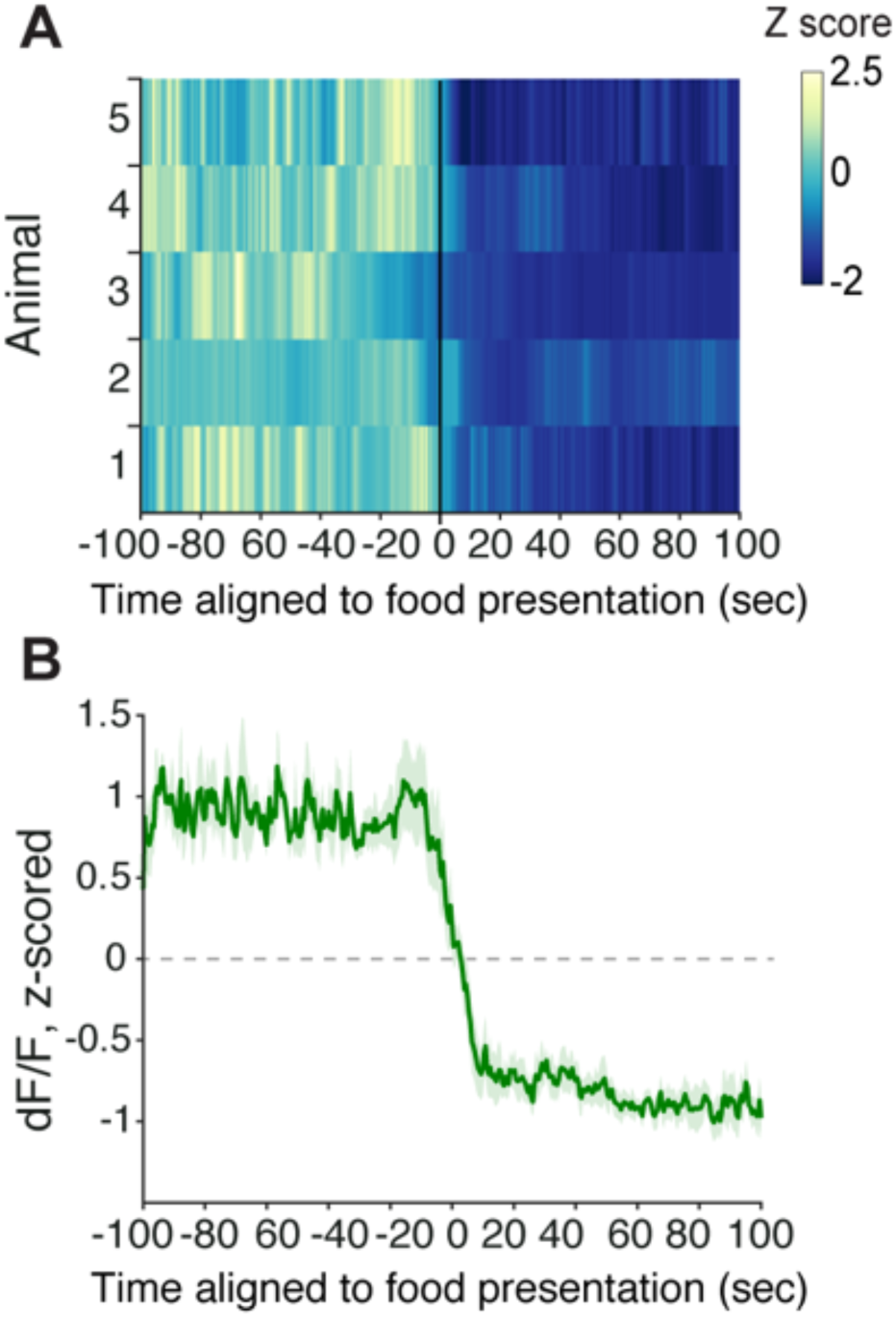
Post-fast refeeding inhibits AgRP neurons. (**A**) Heat map of z-scored dF/F and (**B**) averaged AgRP neuron GCaMP6s activity from individual overnight fasted mice presented with a food pellet, n=5/group, Mean ± SEM.

**Supplemental Figure 5 (Related to Figure 4).**
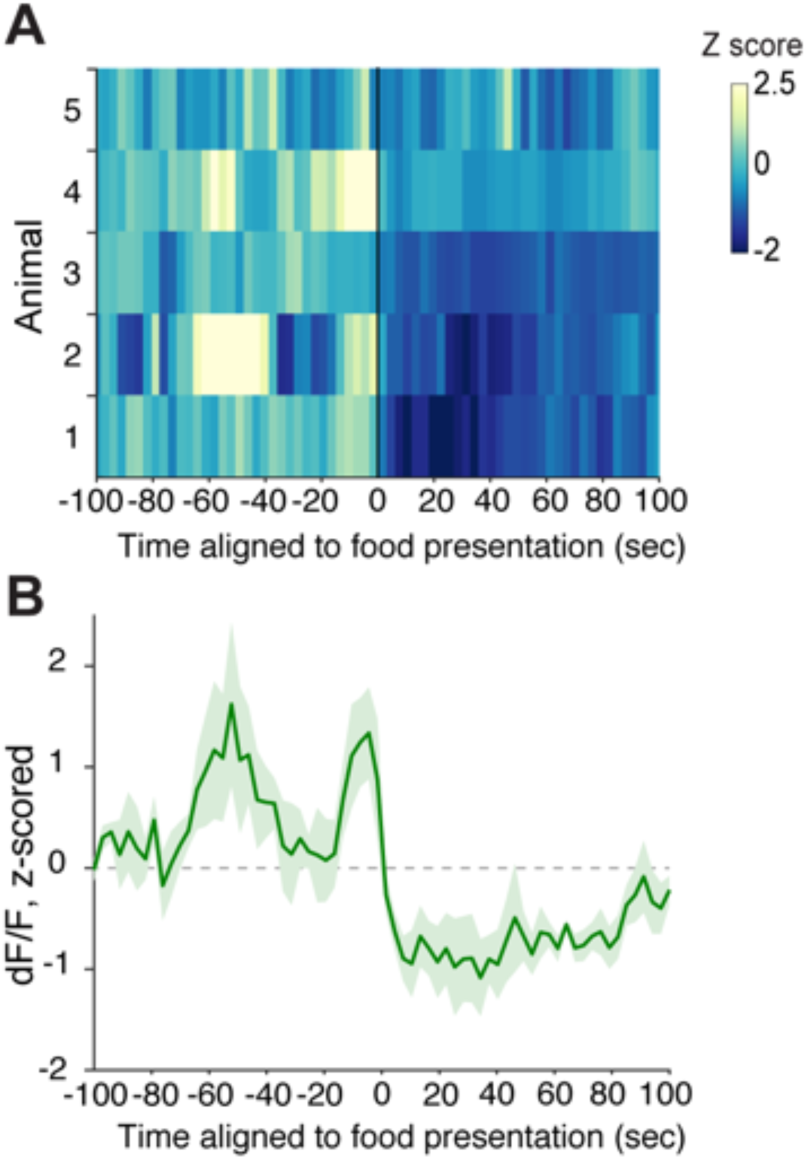
AgRP neuron GCaMP6s activity is reduced by food presentation in ad lib fed mice. (**A**) Heat map of z-scored dF/F aligned to food pellet presentation and (**B**) averaged z-scored GCaMP6s signal. n=5/group, Mean ± SEM.

**Supplemental Figure 6 (Related to Figure 5).**
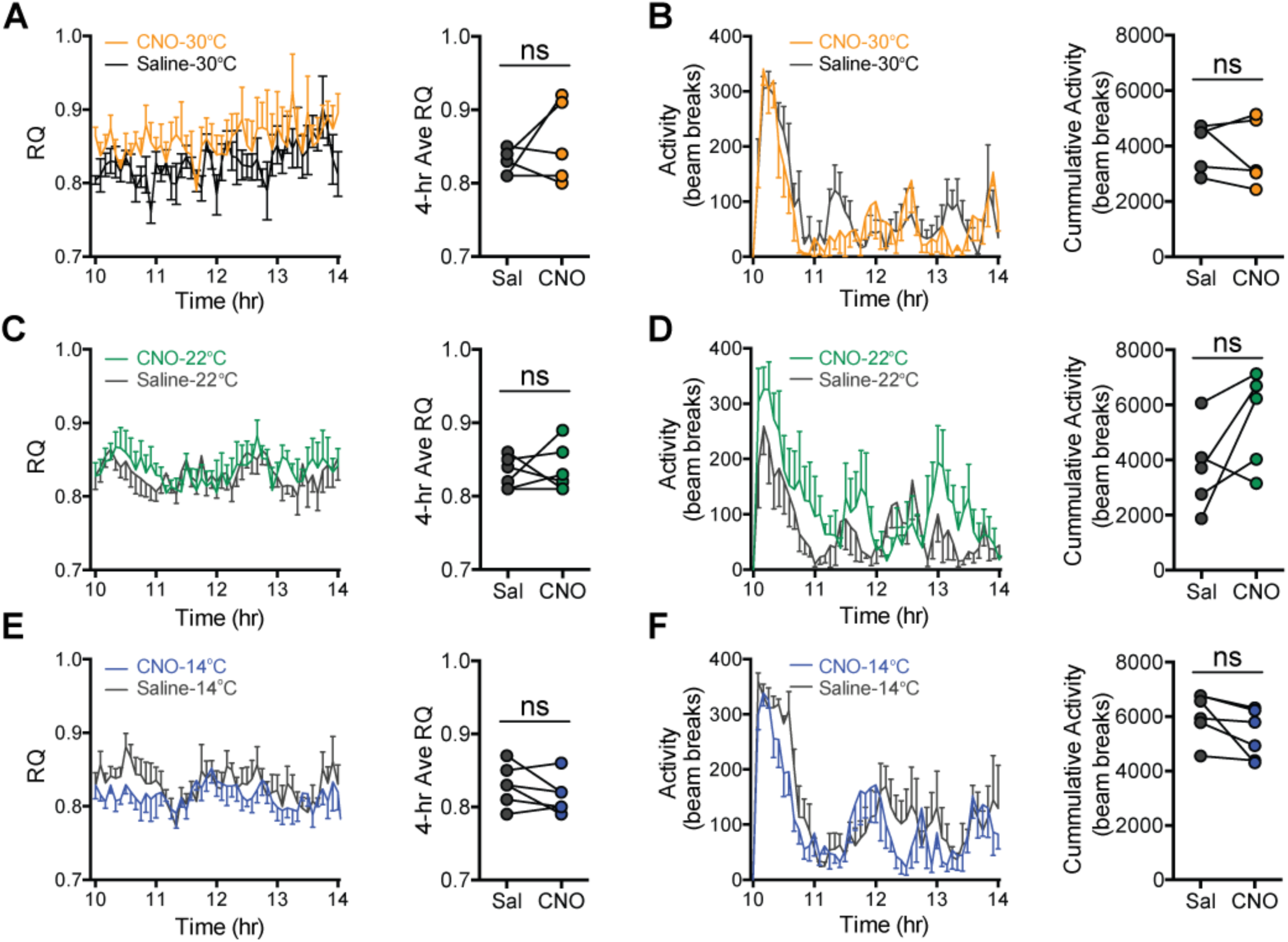
Respiratory quotient and ambulatory activity after AgRP neuron inhibition at three ambient temperatures. (**A, C, E**) 4-hr time-series and mean value for respiratory quotient (RQ), and (**B, D, F**) 4-hr time-series and mean value of cumulative activity in hM4Di-eYFP AgRP-IRES-Cre mice that were housed at either 30°C, 22°C or 14°C after receiving an i.p. injection of either saline or CNO in a randomized, crossover manner, n=6-8/group, Mean ± SEM.

